# Generalization of the Ewens sampling formula to arbitrary fitness landscapes

**DOI:** 10.1101/065011

**Authors:** Pavel Khromov, Constantin D. Malliaris, Alexandre V. Morozov

## Abstract

In considering evolution of transcribed regions, regulatory modules, and other genomic loci of interest, we are often faced with a situation in which the number of allelic states greatly exceeds the population size. In this limit, the population eventually adopts a steady state characterized by mutation-selection-drift balance. Although new alleles continue to be explored through mutation, the statistics of the population, and in particular the probabilities of seeing specific allelic configurations in samples taken from a population, do not change with time. In the absence of selection, probabilities of allelic configurations are given by the Ewens sampling formula, widely used in population genetics to detect deviations from neutrality. Here we develop an extension of this formula to arbitrary, possibly epistatic, fitness landscapes. Although our approach is general, we focus on the class of landscapes in which alleles are grouped into two, three, or several fitness states. This class of landscapes yields sampling probabilities that are computationally more tractable, and can form a basis for the inference of selection signatures from sequence data. We demonstrate that, for a sizeable range of mutation rates and selection coefficients, the steady-state allelic diversity is not neutral. Therefore, it may be used to infer selection coefficients, as well as other key evolutionary parameters, using high-throughput sequencing of evolving populations to collect data on locus polymorphisms. We also carry out numerical investigation of various approximations involved in deriving our sampling formulas, such as the infinite allele limit and the “full connectivity” assumption in which each allele can mutate into any other allele. We find that our theory remains sufficiently accurate even if these assumptions are relaxed. Thus, our framework establishes a theoretical foundation for inferring selection signatures from samples of sequences produced by evolution on epistatic fitness landscapes.

## Introduction

With the advent of high-throughput molecular biology techniques, it has recently become possible to carry out large-scale phenotypic assays in molecular systems. For example, Podgornaia and Laub have mapped all 20^4^ = 1.6 × 10^5^ possible combinations of four key residues in the *E. coli* protein kinase PhoQ, and assayed each mutant for the signaling function mediated by its binding partner PhoP.^1^ The study revealed 1659 functional PhoQ variants, which can be thought of as forming the upper plane on the fitness landscape. The upper plane is divided into several clusters under single-point mutational moves – only sequences within each cluster can mutate into each other without undergoing deleterious moves to the lower plane, where all non-functional sequences reside. This two-plane landscape is highly epistatic – the effect of a given mutation depends on the amino acids at the other three positions, in agreement with previous reports on the primary role of epistasis in molecular evolution.^2,3,4,5^

The picture of fitness landscapes made of multiple interconnected planes is supported by other high-throughput experiments aimed at elucidation of the relationship between gene sequence and function.^3,6,7^ Although these experiments typically yield distributions of mutation fitness effects, the data can often be understood, at least to a first-order approximation, in terms of functional and non-functional sequence variants.^8,9^ Indeed, distributions of fitness effects often appear to be bi-modal in such studies, with a low-fitness peak for strongly deleterious and lethal mutations, and another for weakly deleterious and neutral ones. In comparison to deleterious and neutral mutations, beneficial mutations are relatively infrequent.^6,7,9^ The coarse-graining of the fitness landscape into functional and non-functional states may be refined by introducing additional fitness values, e.g. for weakly deleterious mutations.

Overall, given the astronomically large number of possible sequence variants, we expect the size of neutrally-connected clusters on all fitness planes to be larger than the population size. Then evolutionary dynamics on a multiple-plane landscape will involve periods of neutral search followed by positive selection, in which the bulk of the population moves to a higher-fitness state.^3,10^ Populations evolving on such a landscape will eventually reach a steady state characterized by mutation-drift balance.^11^ This balance determines the statistics of the population, such as the mean and the variance of the number of distinct alleles. Although the population continues to explore new alleles through mutations, the allele statistics do not change anymore once the steady state is reached. In the absence of selection (that is, for evolution on a single neutral plane), the probability of observing a given pattern of allelic diversity in a sample of size n taken from the steady-state population was derived by Ewens.^12^

The Ewens sampling formula can be used to understand the allelic diversity in neutral populations, and test for deviations from the neutral expectation.^13^ However, in order to make quantitative predictions of selection coefficients, it is necessary to extend the Ewens sampling formula to arbitrary fitness landscapes. As noted above, of special interest in molecular evolution are landscapes in which alleles are grouped into two or three distinct fitness states. Such landscapes provide a natural generalization of the completely neutral evolutionary scenario.

Previous work in this area has focused mostly on deriving frequency spectra for either arbitrary fitness landscapes or specific models of selection. In particular, Li obtained the frequency spectrum for a general landscape, and used it to derive expressions for the mean number of alleles in a sample, as well as the mean and the variance of the heterozygosity.^14,15,16^ Ewens and Li derived frequency spectra for landscapes with two and three fitness states, and used them to compute the mean number of distinct alleles and the mean heterozygosity.^17^ Griffiths also derived a general integral expression for the frequency spectrum.^18^

More recently, Ethier and Kurtz have studied allelic diversity in a general model of selection in which fitness of each new allele is completely independent of the fitness of its parent.^19^ This and follow-up work^20,21,22^ has contributed to our understanding of how allelic sampling probabilities are shaped by various forms of selection pressure. In particular, Joyce and Genz have developed effective algorithms for evaluating sampling probabilities.^23^ Finally, Desai et al. have investigated sampling probabilities in a model (previously introduced by Charlesworth et al.^24^ and Hudson and Kaplan^25^) comprised of a sequence of neutral and negatively selected sites.^26^ This model has no epistasis, and therefore can be treated using the Poisson Random Field method.^27^ However, the models of both Ethier and Kurtz and Desai et al. cannot be applied to molecular evolution, which is characterized by prominent epistasis and correlated fitness values.

Here we develop an extension of the Ewens sampling formula to arbitrary fitness landscapes. First, we use the diffusion approximation of population dynamics to derive a general sampling formula valid for any number of alleles *K* (allele *i* is assigned an arbitrary fitness value *f_i_*), mutation rate *μ*, and population size *N*. We assume that the population adopts a steady state characterized by mutation-selection-drift balance. The sampling formula is derived under the assumption that *n* ≪ *N*, where *n* is the total sample size; this assumption is often realistic in populations subjected to high-throughput sequencing.

The most general sampling formula is not amenable to efficient calculations since it involves sums over special functions with a number of terms that increases rapidly with both the sample size and the number of alleles. Therefore, we focus on multiple-plane landscapes applicable in molecular evolution, with only a few (two or more) distinct fitness states. We also make the *N* ≪ *K* approximation, which reflects the fact that the number of possible allelic states in molecular systems is typically much larger than effective population sizes. The resulting sampling formulas are sufficiently tractable to be used to study selection signatures and deviations from neutrality on multiple-plane fitness landscapes for arbitrary mutation rates and selection coefficients. In particular, we study the effective population size approximation^24, 26^ and its limits of applicability. We compare our results with numerical simulations, investigating potential deviations between experiment and theory which may be caused by the differences in evolutionary dynamics between fully connected sequence networks (for which our theory is valid) and more realistic scenarios involving single-point mutations. We also investigate finite network size effects, since our multiple-plane sampling formula is derived in the infinite allele limit.

Our results are applicable to understanding the nature of allelic diversity under selection, mutation and drift, for a vast class of fitness landscapes that are relevant to both molecular evolution and, more generally, evolution in systems where the number of alleles vastly exceeds the population size. Moreover, our sampling formulas form the basis for a quantitative test which can both detect the presence of selection and estimate selection coefficients in epistatic systems under very general and well-defined assumptions. Population-level allele diversity data is increasingly available through high-throughput sequencing techniques, making our approach a practical and timely tool for studying the role played by selection in present-day populations.

## Results

### Steady-state distribution of allele frequencies

We consider a haploid population of fixed size *N*. Each individual in the population has a single allele in the state *i*, with fitness *f_i_*; there are *K* distinct allelic states. Mutations occur at a probability *μ* per generation, replacing the original allele with one of the *K* − 1 remaining alleles. Thus the probability of offspring *A_j_* produced by parent *A*_*i*≠*j*_ is *μ*/(*K* − 1). We can view this system as an “allelic network” with the topology of a complete graph, with *K* vertices representing allelic states and edges representing mutational moves. Stochastic evolution of the population can then be described using Moran^28,29^ or Wright-Fisher^29,30^ population dynamics.

Without loss of generality, we can specify fitness *f_i_* of the allele *A_i_* with respect to an arbitrary reference allele *A_K_*. It is convenient to introduce a *K*-dimensional vector of relative fitnesses multiplied by the population size: 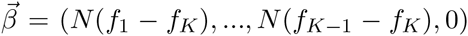. Likewise, we define a *K*-dimensional vector of mutation rates as 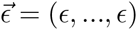, where *ϵ* = *Nμ*/(*K* − 1) for Moran^28,29^ and *ϵ* = 2*Nμ*/(*K* − 1) for Wright-Fisher population dynamics.^29,30^ ^a^ We also introduce *θ* = *Kϵ*, which in the limit of large number of alleles *K* → ∞ becomes *Nμ* and 2*Nμ* for Moran and Wright-Fisher dynamics, respectively. Note that we consider the case of equal mutation rates between alleles, for which the steady state is well-defined.^31^

The evolutionary dynamics of this system in the diffusion limit is described by the forward Kolmogorov equation:

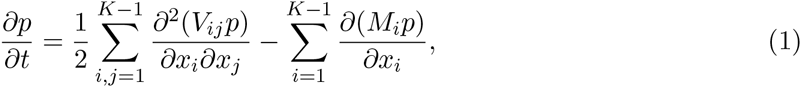

where 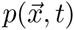 is the joint probability of frequencies of *K* alleles at time 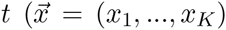 is a *K*-dimensional vector of allele frequencies which occupy a (*K* − 1)-dimensional simplex 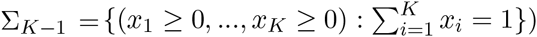, and

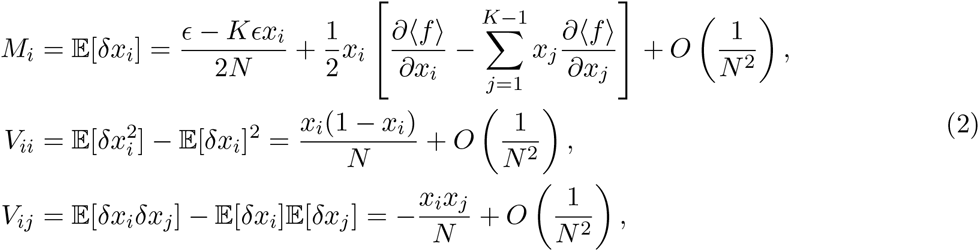

where *x_i_* denotes the frequency of allele *A_i_* in the population, and 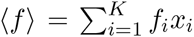 is the mean population fitness.

In steady state ∂*p*/∂*t* = 0, and the distribution of allele frequencies in Eq. 1 is given by^14,15,32^

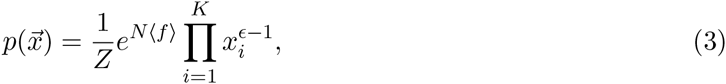

where *Z* is the normalization constant. Eq. 3 can be written more explicitly as

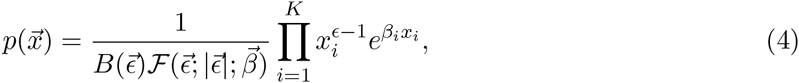

where 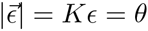 is the *L*_1_-norm of 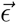,

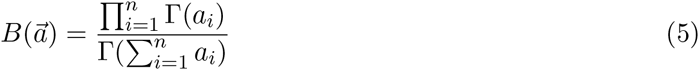

is the generalized beta function written in terms of Gamma functions, and

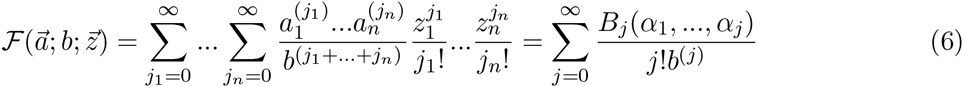

is the confluent hypergeometric function _1_*F*_1_(*a*; *b*; *z*) generalized to vector arguments. Here, *a*^(*j*)^ = Γ(*a* + *j*)/Γ(*a*) is the rising factorial, *B_j_* is the *j*th complete Bell polynomial and 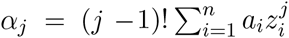.

Differentiation of this function with respect to 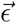 yields

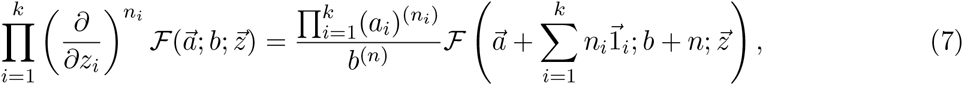

where 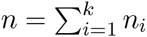 and (1_*i*_)_*j*_ = *δ_ij_*.

### Strongly monomorphic limit

When mutation rate decreases and population size is kept fixed, *ϵ* → 0 and the population becomes monomorphic.^29,33,34,35^ We consider the Fourier transform of the steady-state distribution in Eq. 4:

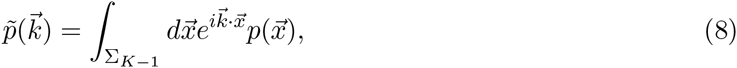

where the integral is over the (*K* − 1)-dimensional simplex. Using Eq. 6, we can write the Fourier transform as a ratio of two generalized hypergeometric functions:

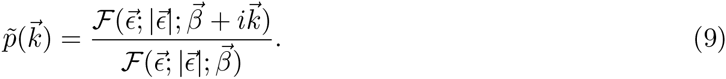

Taking the *ϵ* → 0 limit yields

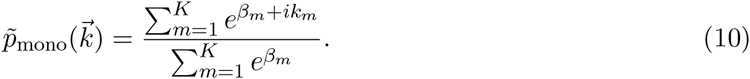

Thus the steady-state distribution in the monomorphic limit is given by:

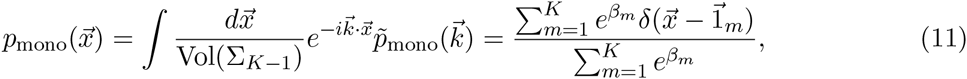

where 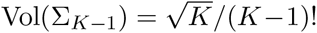 is the volume of the (*K* − 1)-dimensional simplex and (1_*m*_)_*i*_ = *δ_mi_*. Therefore in the *ϵ* → 0 limit the population resides in one of the *K* monomorphic states available to it, with the probability of being in a particular state exponentially weighted by its fitness.^36,37,38^

### Probability of a sample of alleles

Let us now consider a situation relevant to molecular evolution, where the number of alleles *K* is much larger than the population size *N*. In this case, the steady state in terms of allele frequencies is unlikely to be reached on relevant evolutionary time scales. Mathematically, the *K* → ∞ limit of Eq. 4 becomes ill-defined.^39,40^ Nonetheless, the steady state is well-defined in terms of allelic *counts* rather than frequencies of specific alleles.^11^ In other words, the allelic diversity of the population (e.g. as characterized by the mean and the variance of the distribution of the number of distinct allelic types) is tractable and will no longer change in steady state, although new alleles will continue being explored through mutation.

One is often interested in statistical properties of a sample of alleles of size *n* ≪ *N* obtained from the population. Let us consider a simple example of a population evolving on a small allelic network with *K* = 5 allelic types *A, B, C, D, E*. Suppose that in sampling *n* = 4 alleles from the population we first observe allele *A*, then *C*, then *A* again, and finally *D*. We can record this sequence of alleles as an ordered list (*A, C, A, D*). However, typically we are not interested in the order in which alleles appear in the sample, and therefore record the result as an unordered list {*A, A, C, D*}, which shows that allele *A* has appeared twice and alleles *C* and *D* have appeared once each. ^b^ Alternatively, we can record non-zero allelic counts, which gives us *n_A_* = 2, *n_C_* = 1, *n_D_* = 1. Finally, we can dispense with the allele labels altogether, identifying each allele in the sample as either new or already seen. In this case, we are left with an unordered list of counts {2, 1, 1}, meaning that we have observed 4 alleles of 3 different types, with one type represented by two alleles and the other two types by one each. In general, we will refer to {*n*_1_, …, *n_k_*} as the (unordered) allelic counts. The allelic counts can also be summarized in terms of a histogram which records how many groups of *j* identical alleles occur in the sample, with *j* ranging from 1 to *n*. In our example, there is one group of two identical alleles and two groups of one allele each, so that (*A, C, A, D*) is recorded as the allelic histogram (*a*_1_ = 2, *a*_2_ = 1, *a*_3_ = 0, *a*_4_ = 0). ^c^

We now derive the probability 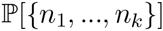 of observing an unordered sample {*n*_1_, …, *n_k_*}, given that the population has reached steady state in terms of its allelic diversity. Before treating the general case, we illustrate our approach using a toy example with *K* = 3 allelic types: 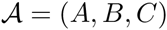. We wish to calculate the probability of observing the {2, 1} unordered configuration in a sample of size *n* = 3, which is assumed to be much less than the population size *N*. There are 18 ordered configurations that contribute to this probability:

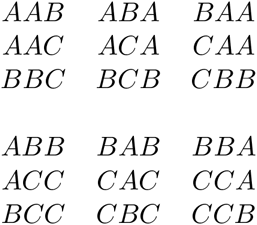

In particular, the probability of choosing *A* first, then *A* again and finally *B* is

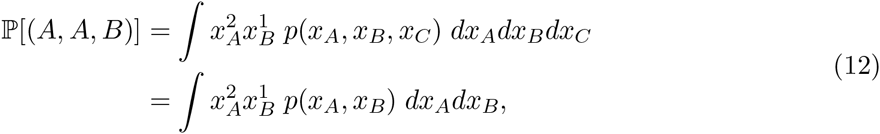

where *p*(*x_A_, x_B_, x_C_*) is given by Eq. 4. Consequently, the probability of observing two *A*’s and one *B* in *any* order is given by^41^

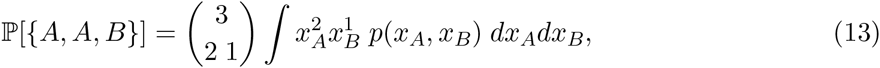

where 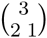 is the multinomial coefficient. Introducing a set 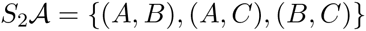, which permutes allelic identities in an ordered manner (i.e., the overall allele ordering from *A* to *B* to *C* is preserved in each pair of alleles), we can take into account the first 9 configurations in the table above:

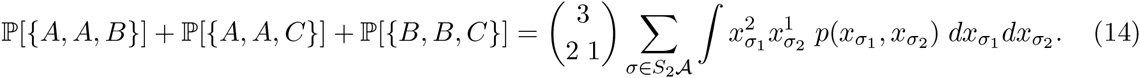

In order to include the last 9 configurations in the table, we need to switch the order of the alleles: {(*A, B*), (*A, C*), (*B, C*)} → {(*B, A*), (*C, A*), (*C, B*)}. But switching the alleles in each pair amounts to replacing 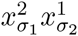 with 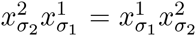 in Eq. 14. Thus we can summarize the entire table by introducing a set 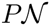 of all distinct permutations of allelic counts 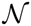, which determine the powers to which the allelic frequencies are raised in Eq. 14. In our example 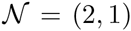 and 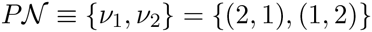. Therefore,

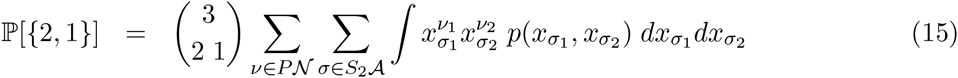

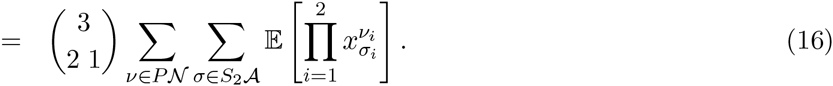

The above example can be easily generalized to describe the probability 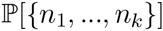 of observing an unordered list of counts, {*n*_1_, …, *n_k_*}. Note that the sample size is 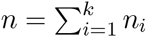, and that *k* distinct allelic types are observed. First, we enumerate all *K* alleles, forming a unique ordered list 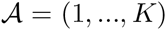. Second, we choose a subset *σ* = (*σ*_1_, …, *σ_k_*) of size *k* from 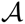 without replacement, so that the allelic order is preserved: *σ*_1_ < … < *σ_k_* (note that no subsets are allowed to contain repeating elements of 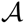). Then 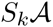 can be naturally defined as a set which contains all ordered subsets of 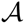 of size *k*. Finally, as before 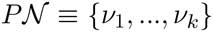 is a set of all distinct permutations of allelic counts 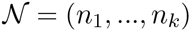. With these definitions,

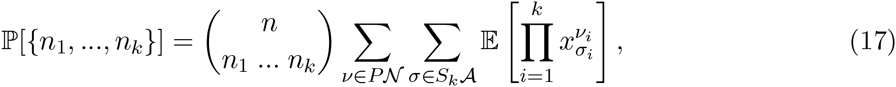

where 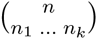 is the multinomial coefficient, and the expectation is calculated with respect to the steady-state allele distribution, Eq. 4.

We can use the probability distribution over unordered configurations (Eq. 17) to compute the distribution of the number of different allelic types *k*:

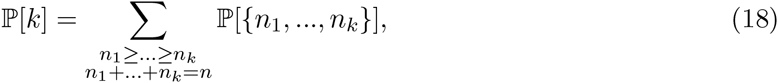

where the summation runs over all ordered partitions of *n* into *k* positive integers.

### Sampling formula for the arbitrary fitness landscape

As Eq. 17 demonstrates, evaluation of sample probabilities requires calculation of moments of allele frequency distributions. This is most easily accomplished by taking derivatives of the normalization constant 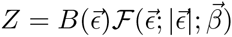 with respect to the components of 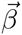:

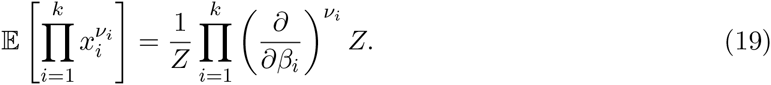

Using Eq. 7, we obtain:

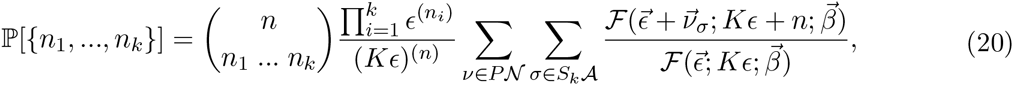

where equal mutation rates are assumed for all alleles and 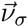 is a *K*-dimensional vector whose *σ_i_*-th components are *ν_i_* (*i* = 1, …, *k*) and all the other components are zero. As discussed above, the sum over *σ* extends over all distinct subsets of *k* alleles sampled from *K* uniquely ordered alleles and subject to the *σ*_1_ < … < *σ_k_* constraint. Therefore 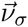 has *K* − *k* zero and *k* non-zero components which are distributed according to *σ*. The sum over *ν* extends over all distinct permutations of allelic counts which sum up to *n*. Eq. 20 is valid for arbitrary fitness landscapes and arbitrary *K*.

### Neutral limit of the sampling formula

When all alleles have the same fitness, the general sampling formula given by Eq. 20 should reduce to the Ewens formula for neutral evolutionary dynamics.^11,12^ Indeed, with all *β_i_* set to zero, the generalized hypergeometric function 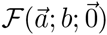 (Eq. 6) becomes 1. Then for the finite number of alleles *K*

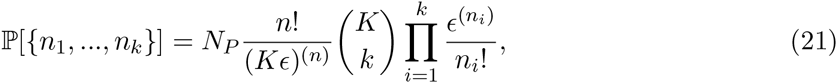

where 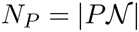 is the total number of distinct permutations of allelic counts (*n*_1_, …, *n_k_*). In the limit of an infinite number of alleles *K* → ∞, Eq. 21 becomes

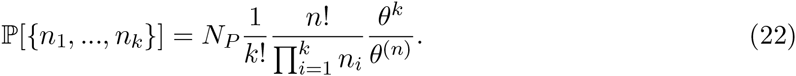

Changing variables to allelic histogram counts yields 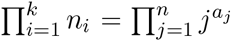 and 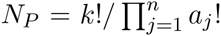, resulting in

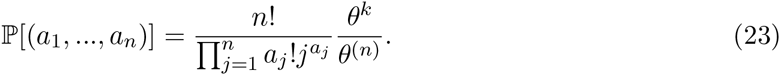

Eq. 23 is known as the Ewens sampling formula.^11,12^

### Sampling formula for populations with two fitness states

As a straightforward generalization of the neutral case, consider a system with *I* alleles of fitness *f*_1_ and *K* − *I* alleles with fitness *f*_2_ = *f*_1_. Thus the fitness landscape consists of two interconnected “planes”. We can assume without loss of generality that alleles 1, …, *I* belong to the first plane and alleles *I* + 1, …, *K* belong to the second plane. Then *γ* = *I*/*K* defines a fraction of nodes on the first plane, and the fitness vector is 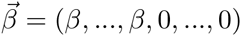 with *I* non-zero entries followed by *K* − *I* zeros. Then it can be shown that for finite *K* the sampling probability is given by (Appendix A):

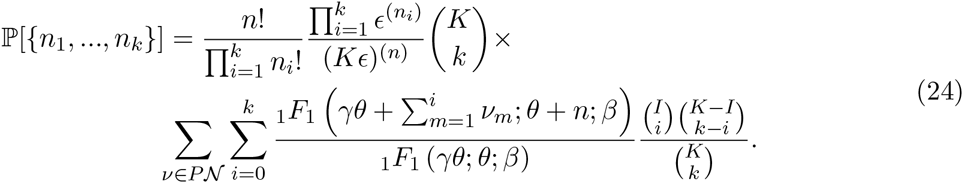

Taking the infinite allele (*K* → ∞) limit with *γ* fixed, we arrive at

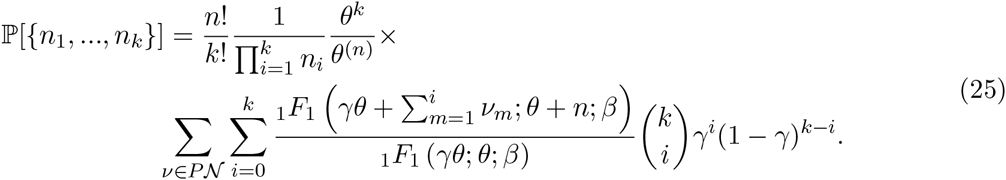

Thus hypergeometric sampling of Eq. 24 reduces to binomial sampling in the infinite allele limit.

### Sampling formula for populations with multiple fitness states

Let us now generalize the result of the previous section to the case of multiple fitness states: each allele can be assigned a distinct fitness value *f_m_, m* = 1, …, *M*. In other words, the fitness landscape consists of multiple planes, with *I_m_* = *γ_m_K* nodes of fitness *f_m_* on the *m*th plane, so that 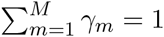. Then the sampling probability for finite *K* is given by

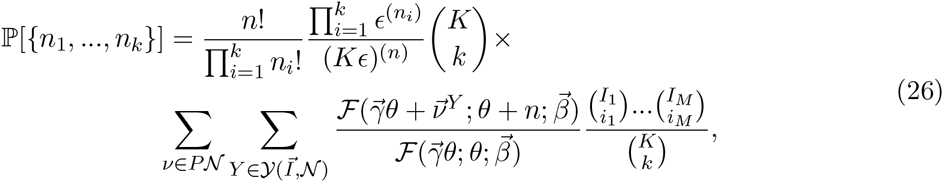

and its infinite allele limit is given by

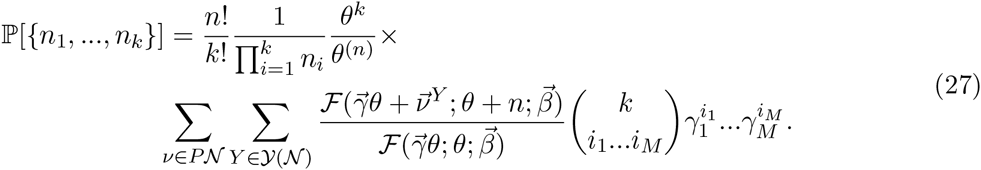

The sums in Eqs. 26 and 27 take into account all possible ways of sampling *n* alleles from *M* planes (Fig. 1). To explain these sums, let us imagine distributing *n* books over *M* shelves. The books come in *k* indivisible volume sets, and the *i*th set has *ν_i_* identical books in it. We would like to find all book-to-shelf arrangements, keeping in mind that shelves have finite capacities: only *I_m_* books can be placed on the mth shelf. One way to describe any book-to-shelf arrangement is to use an *M*-dimensional vector 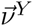 which records how many books are placed on each shelf. For example, if *M* > *k*, a vector 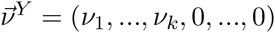 with *M* − *k* zeros following *k* non-zero entries describes placing volume sets on shelves in a particular order: the first volume set goes on the first shelf, the second volume on the second shelf and so on (assuming that the shelves are large enough to accommodate the volume sets), until no more books are left, so that the remaining *M* − *k* shelves remain empty. Permutations of this arrangement, expressed as permutations of 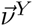 vector elements, are also allowed (again, assuming that all the shelves are large enough). We can also put more than one volume set on a single shelf, leading to arrangements such as (*ν*_1_ + *ν*_2_, *ν*_3_, …, *ν_k_*, 0, …, 0) with *M* − *k* + 1 zero and *k* − 1 non-zero entries. As before, this arrangement is allowed only if the number of books on each shelf does not exceed shelf capacities. Note that the question of capacity does not arise in the infinite allele limit, since the shelves become effectively infinitely long.

**Figure 1:**
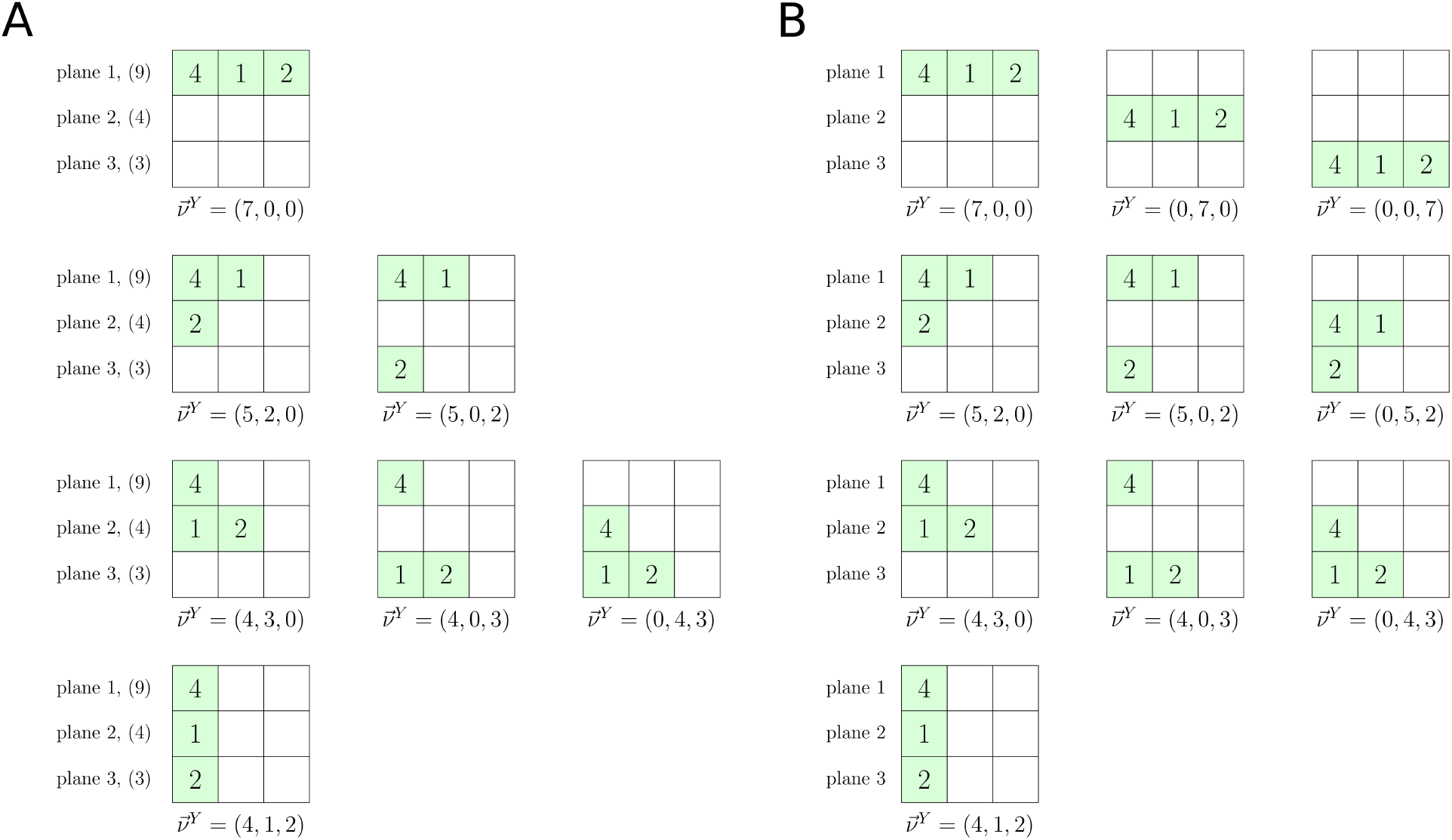
Illustration of summations over 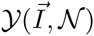 and 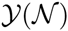 in Eqs. 26 and 27 respectively, for a list of allelic counts 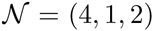. (A) The finite plane case. Finite plane capacities are shown in parentheses. (B) The infinite plane case.

In order to systematically list all the arrangements for volume sets (*ν*_1_,…, *ν_k_*), we follow a simple rule: if the *k*th set of *ν_k_* books is placed on the *m*th shelf, the (*k* + 1)th set of *ν*_*k*+1_ books goes either on the same shelf or on the *m*′th shelf with *m*′ > *m*. Taking elements of (*ν*_1_, …, *ν_k_*) one by one and changing the initial shelf (onto which the 1st volume set is placed) and the number of volume sets on each shelf, we can generate a set of all permutations of 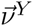 elements. We shall call this set 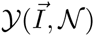 since it depends on both the shelf capacities 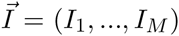 and the volume sets 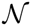. In the limit of infinite shelf capacity the dependence on shelf sizes disappears, and the set of all permutations will be called 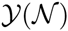. To include all possible arrangements, we need to perform the book-placing procedure for each distinct permutation of 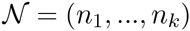.

Now, if we replace shelves with fitness planes and volume sets with allelic counts, we obtain an algorithm for generating all allowed placements of allelic counts on fitness planes. The non-negative indices *i*_1_ … *i_M_* in Eqs. 26 and 27 represent the number of volume sets (allelic counts) on each shelf (fitness plane). The distribution of alleles among fitness planes of finite capacity is illustrated in Fig. 1A for *M* = 3 and a vector of allelic counts 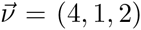; the infinite-plane case is shown in Fig. 1B.

Next, let us consider the monomorphic limit of Eq. 27: *θ* → 0 with finite *β* and *γ*. It can be shown that

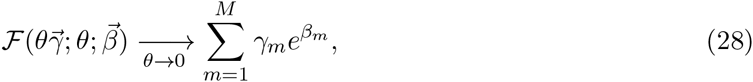

leading to

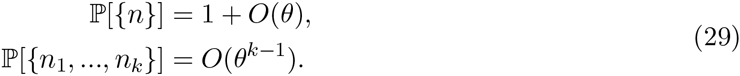

Therefore, as expected, the 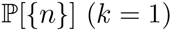 term predominates in the monomorphic limit.

By construction, Eqs. 25 and 27 reduce to the neutral limit (Eq. 22) when all fitness values are the same. In addition, the neutral limit is reproduced in the strongly polymorphic, mutation-dominated limit, defined as *θ* → ∞ with finite *β* and *γ*. In this limit,

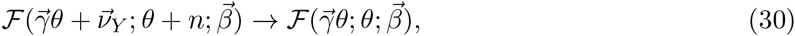

and Eq. 27 reduces to the neutral result. This is expected since selection effects become vanishingly small in this regime.

### Frequency spectrum for arbitrary fitness landscapes

The frequency spectrum Φ(*x*) is a standard way of characterizing allele frequency distributions in evolving populations;^42^ Φ(*x*)*dx* is defined as the number of alleles in the population with frequency in the (*x, x* + *dx*) range. Therefore, according to Eq. 4 the steady-state frequency spectrum is given by

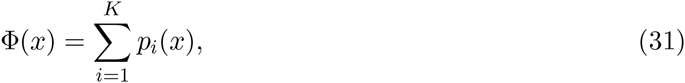

where 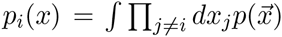 is the marginalized allele frequency distribution for the *i*th allele. Note that according to Eq. 31 Φ(*x*) is normalized as follows:

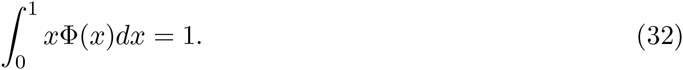

Correspondingly, *x*Φ(*x*)*dx* is the probability that an allele randomly drawn from the population has its frequency in the population in the (*x, x* + *dx*) range. The frequency spectrum can be used to find 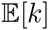, the mean number of distinct alleles in a sample of size *n*:

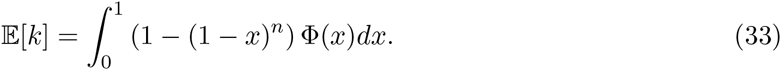

For a landscape with *M* distinct fitness values, the frequency spectrum is given by (Appendix B)

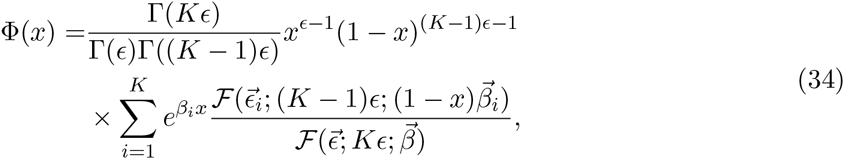

where the (*K* − 1)-dimensional vectors 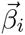 and 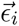 are obtained from *K*-dimensional vectors 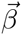 and 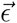 by removing their *i*th components. The formula above is valid for arbitrary number of alleles *K*, mutation rate, and population size.

Using Eq. 34, the expected number of distinct alleles in a sample of size *n* can be computed as

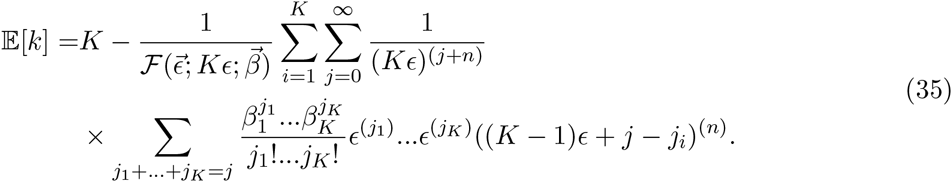

In the *K* → ∞ limit (and assuming that all *M* fitness planes also become infinite in this limit), Eq. 34 simplifies to

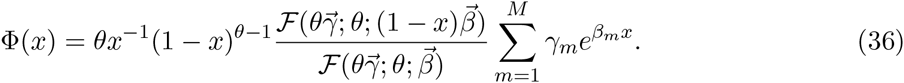

Furthermore, in the case of two fitness states (*M* = 2) we can simplify Eq. 36 using Eq. 55:^17^

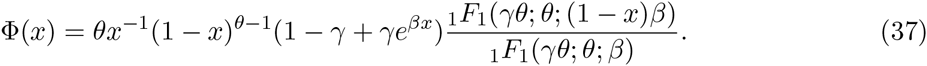

For neutral evolution, we set all *β_i_* to 0; Eq. 36 then yields

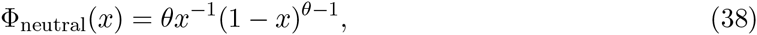

which looks like two-allele steady-state allele frequency distribution. In the strongly monomorphic limit and the absence of selection, the steady-state distribution (Eq. 4) simplifies to (Eq. 11):

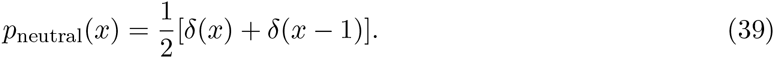

Since Φ(*x*) = *Kp*(*x*), we obtain

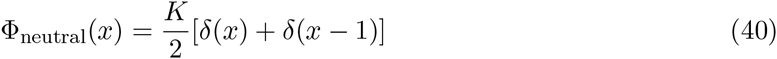

in the monomorphic limit of neutral evolution.

Using Eq. 33, we can obtain a standard expression for the mean number of alleles observed in neutral evolution:^11,15,17^

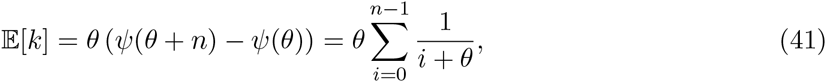

where *ψ*(*z*) = Γ′(*z*)/Γ(*z*) is the digamma function. Note that 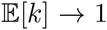 in the monomorphic limit (*θ* → 0) and 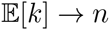 in the strongly polymorphic limit (*θ* → ∞).

Finally, we observe that Eq. 41 can also be derived by setting all *β_i_* = 0 in Eq. 35:

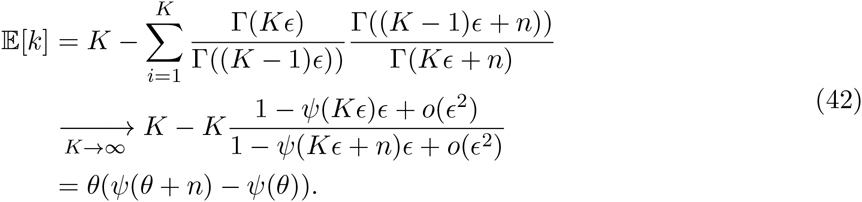

In the monomorphic limit (*θ* → 0), Eq. 36 becomes

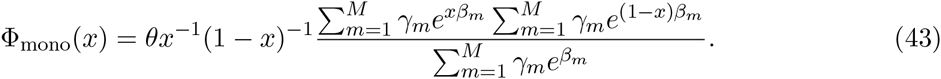

In this limit, *x*Φ_mono_(*x*) is non-zero only if *x* ≃ 1, where Φ_mono_(*x*) ≃ *θx*^−1^(1 − *x*)^−1^. Note that selection effects disappear: the entire population is in the same allelic state due to genetic drift, performing a random walk on the uppermost fitness plane.

In the strongly polymorphic, mutation-dominated limit (*θ* → ∞), Eq. 36 simplifies to

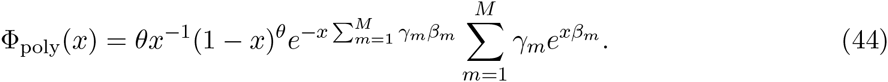

In this case, *x*Φ_poly_(*x*) is peaked around *x* ≃ 0, with Φ_poly_(*x*) ≃ *θx*^−1^(1 − *x*)^*θ*^. The population is completely delocalized, with each member of the population in a distinct allelic state and negligible selection effects. This is not surprising since mutation dominates both selection and genetic drift in this limit.

### Fitness landscape models

We have carried out validation of our theoretical predictions against numerical simulations. We have used the Moran model of population genetics^11,28^ to evolve a population of *N* = 10^3^ haploid organisms, each of which could be in one of *K* allelic states. Specifically, at each step a parent is chosen by randomly sampling the population with weights proportional to the fitness of each individual. An offspring is then produced as an exact copy of the parent. Next, the offspring undergoes mutation with the probability *μ*. Finally, the population is uniformly sampled to choose an organism that will be replaced by the offspring, keeping the overall population size constant. Probabilities of sampling *n* individuals from the population were calculated as averages over 10^6^ samples gathered from 10^3^ independent runs. For each run, a randomly generated initial population was evolved to steady state, after which *n* individuals were sampled from the population with replacement 10^3^ times, waiting ~ 1/*μ* generations between samples.

We consider two types of models with different mutational moves. In the first model, each allele is allowed to mutate into any of the other *K* − 1 alleles with equal probabilities. We call this model fully-connected (FC); it corresponds to our theory which was developed for FC networks. In the second model, a more realistic move set of single-point mutations is implemented: each organism’s genome is represented by a sequence of integers *a*_1_…*a_L_* of length *L*, where 0 ≤ *a_i_* ≤ *A* − 1. A mutation replaces an integer at a randomly chosen site with one of the remaining *A* − 1 integers; all the replacements have equal probabilities. We call this model a single-point mutation (SPM) model.

Finally, we assign a fitness value to each allele. We focus on the landscapes in which alleles can have either low or high fitness values (the “two-plane” model), or low, intermediate, and high fitness values (the “three-plane” model). The fractions of alleles found in each plane are given by 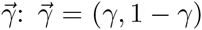 for the two-plane model and 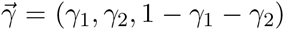 for the three-plane model. In the FC model, the mutational neighborhood of each allele is the same, so that any desired allele fractions 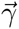 can be implemented. However, in the SPM model the fractions of neutral, beneficial and deleterious moves in each plane will depend on 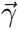 and the assignment of states to planes. We wished to produce non-trivial distributions of neutral moves on the fitness planes, with mutational neighborhoods of some alleles being completely neutral in each plane. Another condition was that the number of alleles in each plane should decrease with its fitness, to reflect the fact that higher-fitness solutions are harder to find.

To fulfill these requirements, we chose to assign fitness values in the SPM model in the following way. We use the sequence length *L* = 10 and the alphabet size *A* = 4. For each sequence *a*_1_…*a_L_* we compute a score *z* = *a*_1_ + … + *a_L_*. We compare these scores with a set of cutoffs (*c*_1_, …, *c*_*M*−1_) for the *M*-plane landscape. For the two-plane landscape, the fitness is 1 if *z* ≤ *c*_1_, and 1 + *s* otherwise. We use the cutoff *c*_1_ = 17, which yields 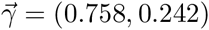. For the three-plane landscape, if *z* ≤ *c*_1_ the fitness is 1, if *c*_1_ < *z* ≤ *c*_2_ the fitness is 1 + *s* − Δ*s*, and if *z* > *c*_2_ the fitness is 1 + *s* + Δ*s*. We choose the cutoffs *c*_1_ = 17 and *c*_2_ = 21, which lead to 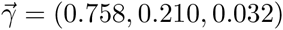. In order to compare FC and SPM simulations directly, we use the same values of 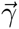 in the corresponding FC models.

Note that in the neutral case the exact mapping between *θ* and *μ* is given by *θ* = *Nμ*/(1 − *μ*) for the Moran model.^11^ However, it is unclear if this mapping can be extended to the non-neutral cases considered here. In any event, for the population size and the values of *θ* investigated below, *μ* = *θ*/(*N* + *θ*) ≃ *θ*/*N*. Therefore, we use the diffusion theory result *θ* = *Nμ* in comparing theoretical predictions with numerical simulations.

### The effective population size approximation

In the monomorphic limit, we expect the effective population size (EPS) approximation to hold:^24,26^ population dynamics is neutral but with the rescaled population size *N*\*. Indeed, in the two-plane case Eq. 25 reduces to

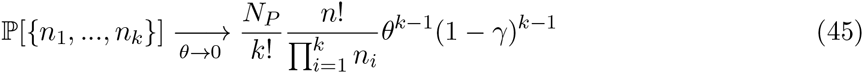

in the *θ* → 0 limit, which corresponds to the *s* ≫ *μ* regime when *β* is finite; Eq. 45 is the same as the neutral sampling formula (Eq. 22) in the monomorphic limit if the population size is rescaled: *N* → *N*\* = (1 − *γ*)*N*. This result can be generalized to the landscape with multiple fitness planes, in which case

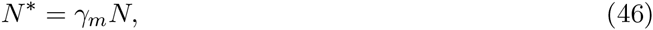

where *γ_m_* is a fraction of nodes with the highest fitness.

However, the EPS approximation breaks down in the polymorphic regime. Indeed, if we take the *N* → ∞ limit, which keeps *β*/*θ* finite (i.e., the ratio of selection and mutation forces remains finite as population size increases), it can be shown for the two-plane landscape that

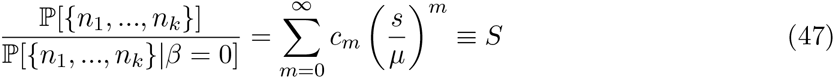

where 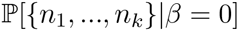 is given by Eq. 22, and coefficients *c_m_* depend solely on *n*_1_, …, *n_k_*. Since the right-hand side of Eq. 47 does not depend on the population size, it can be used to define *N*\* = *S*^1/(*k−n*)^*N*. However, this definition will be sample-specific, due to dependence of *S*^1(*k−n*)^ on *n*_1_, …, *n_k_*. Thus there is no global rescaling of the population size in the strongly polymorphic regime, and evolutionary dynamics is non-neutral.^26^

### Detection of selection signatures

As discussed above, in general we expect allele diversity to deviate from neutrality, making it possible to detect selection signatures using sequences sampled from a population as input. To investigate non-neutral population dynamics, we compute probabilities for all partitions {*n*_1_, …, *n_k_*} of n alleles sampled from the population evolving under selection, and compare them with steady-state partition probabilities obtained under neutral evolution and the EPS approximation.

We use the Kullback-Leibler (KL) divergence to quantify the difference between two probability distributions:^43^

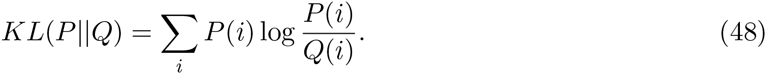

For the two-plane system, we first compare partition probabilities under selection, 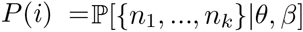, with the corresponding neutral probabilities, 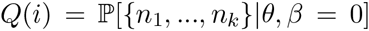. Here, *i* labels distinct partitions. In Fig. 2A, we plot the KL divergence as a function of the mutation rate and the selection strength for the two-plane fitness landscape. We observe that evolutionary dynamics is essentially neutral if selection is weak (*s* ≤ *μ*); in addition, the range of selection coefficients for which neutrality holds increases in the monomorphic regime (*Nμ* ≤ 1). On the other hand, population statistics is clearly non-neutral when the population is polymorphic and the separation between the two fitness planes is large. Next, we compute the KL divergence *KL*(*P*||*Q*\*) between the EPS probability distribution, 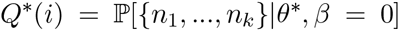, where *θ*\* = (1−*γ*)*θ*, and *P*(*i*) (Fig. 2B). We see that the EPS approximation fails in the polymorphic, weak-selection regime. Overall, the neutral and EPS approximations are approximately complementary – for example, in the strong-selection (*s* ≫ *μ*), polymorphic regime, when evolutionary dynamics becomes non-neutral, it is well approximated by the EPS model.

**Figure 2:**
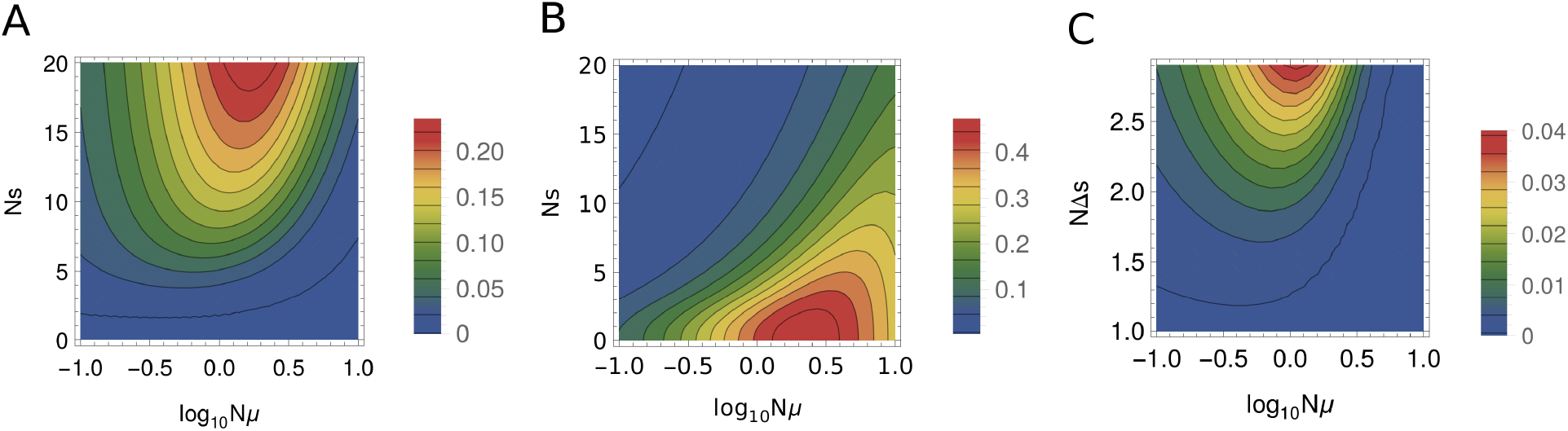
Probabilities of all possible partitions of *n* = 3 alleles ({3}, {2, 1} {1, 1, 1}) sampled from the population of size *N* = 10^3^. (A) and (B) KL divergences for the two-plane fitness landscape as a function of the mutation rate *Nμ* and the selection coefficient *Ns* scaled by the population size, for partition probabilities with and without selection (A), and partition probabilities with selection compared with the EPS approximation (Eq. 45) (B). (C) KL divergences for the sampling probabilities of all possible partitions on a three-plane vs. two-plane landscape. Alleles in the three planes have fitnesses 1, 1 + *s* − Δ*s* and 1 + *s* − Δ*s* respectively, with *Ns* = 6 for both two and three-plane landscapes.

In Fig. 2C we show KL divergences between partition probability distributions on two- and three-plane fitness landscapes. We observe that the partition probabilities are essentially two-plane (i.e., there are no selection signatures indicating presence of intermediate-fitness alleles) if the population is monomorphic (*Nμ* ≤ 1), or if the distance between the two upper planes is smaller than the mutation rate (Δ*s* ≤ *μ*). However, there is a considerable parameter region in which deviations between two and three-plane sampling probabilities appear to be significant (with KL divergences between the two distributions of 0.01 or more), making it possible to detect three distinct fitness states in the sampling data.

### Mutation load

By definition, the mutation load is given by^29,35^

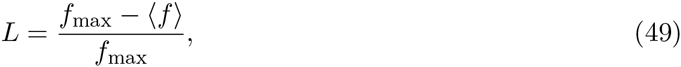

where *f_max_* is the maximum fitness, and 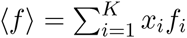 is the mean population fitness. To estimate the mutation load at steady state, we compute the expected value of the mean population fitness over multiple realizations of the stochastic process:

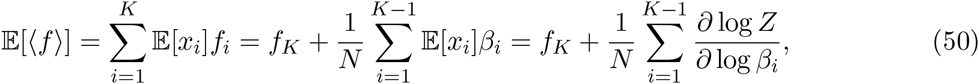

where 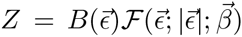 is the normalization in the steady-state allele frequency distribution (Eq. 4). Choosing *f_K_* to be the maximum fitness, we obtain the following expression for the mutation load:

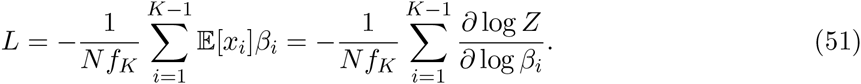

For the two-plane system, Eq. 51 yields (Appendix A):

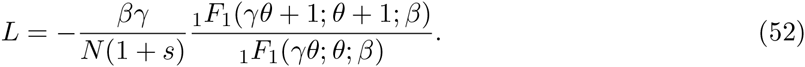

Note that in the two-plane system

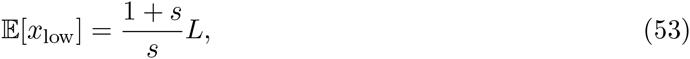

where 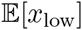 is the average fraction of the population on the lower plane.

Mutation loads for the two-plane fitness landscape are shown in Fig. 3A over a range of selection strengths and mutation rates. As expected, we observe that the largest deviations from the maximum fitness occur in the strong-mutation, strong-selection regime, where a fraction of the population is constantly displaced to the lower plane by mutation, incurring a fitness cost. Correspondingly, at a given value of selection strength the mutation load increases with the mutation rate. In the monomorphic regime the mutation load is vanishingly low because the entire population condenses to a single allelic state and moves randomly on the upper plane. The fraction of the population on the lower fitness plane is shown in Fig. 3B. The fraction is high when the separation between the two planes is low and, at a fixed separation, it increases with the mutation rate.

**Figure 3:**
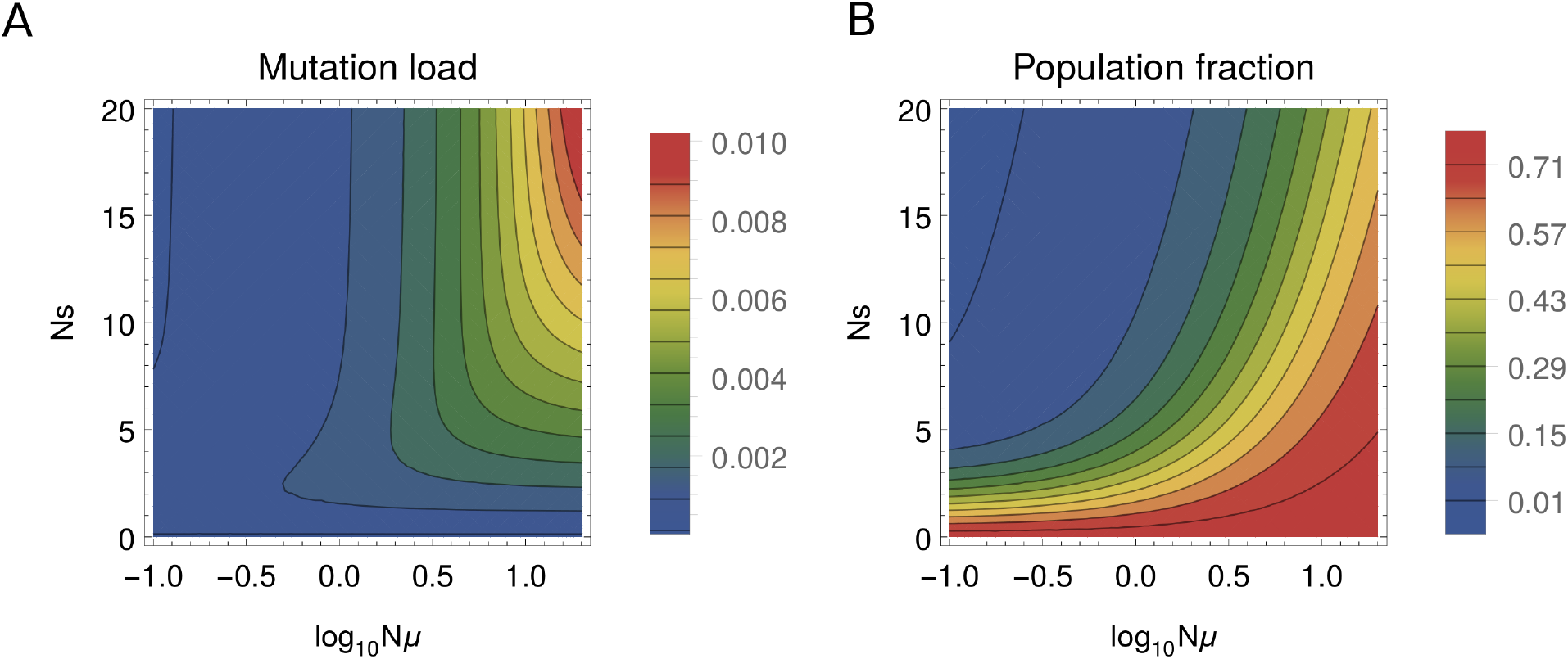
(A) Mutation load (Eq. 52) and (B) the population fraction on the lower plane (Eq. 53) for the two-plane fitness landscape, as a function of the mutation rate (*Nμ*) and the selection strength (*Ns*) rescaled by the population size.

### Partition probabilities on fully-connected vs. single-point-mutant networks

Our theoretical results have been developed for fully-connected networks in which an allele can mutate into any other allele. However, this model is not realistic for protein or nucleotide sequences, in which mutational neighborhoods of a given sequence consist of single-point mutants, i.e. sequences that differ from each other at only a single site. Here we investigate how partition probabilities change if we switch from the FC to the SPM allele network described above. In Fig. 4, we compare theoretical predictions with numerical simulations on the FC and SPM networks in the two-plane system. Overall, we observe excellent agreement between theory and simulations on FC networks. Furthermore, we see that the agreement between SPM simulations and our theoretical results is reasonable: in nearly all cases, the predicted ranking of the sample partitions, as well as the ranking within any given sample partition with respect to *Ns*, are preserved. The largest discrepancies occur in the weakly polymorphic (*Nμ* = 1), strong-selection regime (*Ns* = 6, 13).

**Figure 4:**
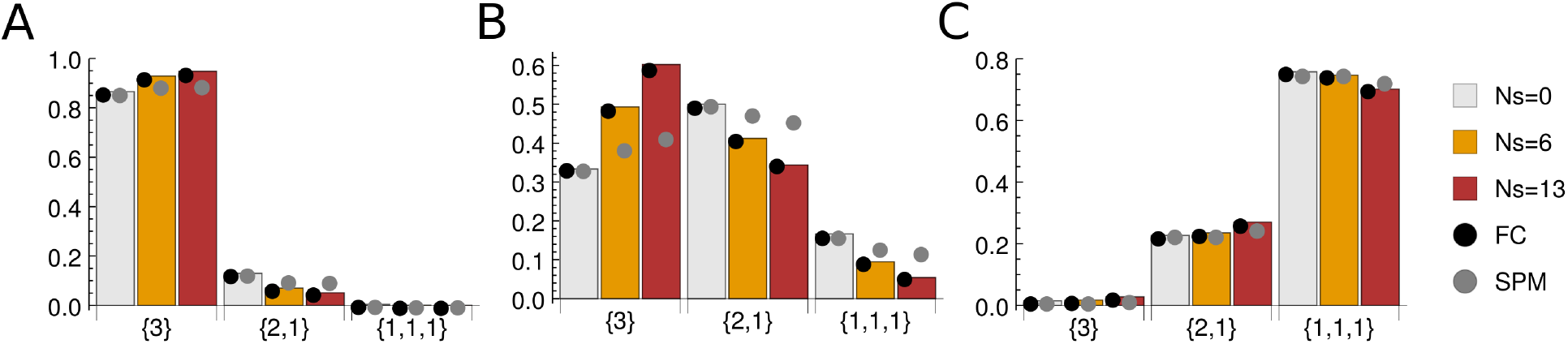
Partition probabilities for the two-plane fitness landscape. Shown are sampling probabilities of all partitions with *n* = 3: {3}, {2, 1}, {1, 1, 1}. Bars: theoretical predictions in the infinite allele limit. Black circles: numerical simulations on the FC sequence network. Grey circles: numerical simulations on the SPM sequence network. In all simulations, alphabet size *A* = 4, sequence length *L* = 10, and population size *N* = 10^3^ were used. Partition probabilities were estimated from 10^6^ samples as described in the main text. (A) Monomorphic population, *Nμ* = 0.1. (B) Weakly polymorphic population, *Nμ* = 1.0. (C) Strongly polymorphic population, *Nμ* = 10.0. Note that the corresponding KL divergences are listed in Table 1.

The situation is qualitatively similar when a three-plane fitness landscape is considered (Fig. 5). We again observe excellent agreement between theory and FC simulations and, overall, reasonable agreement between theory and SPM simulations, with the largest discrepancies again occurring in the weakly polymorphic, strong-selection regime.

**Figure 5:**
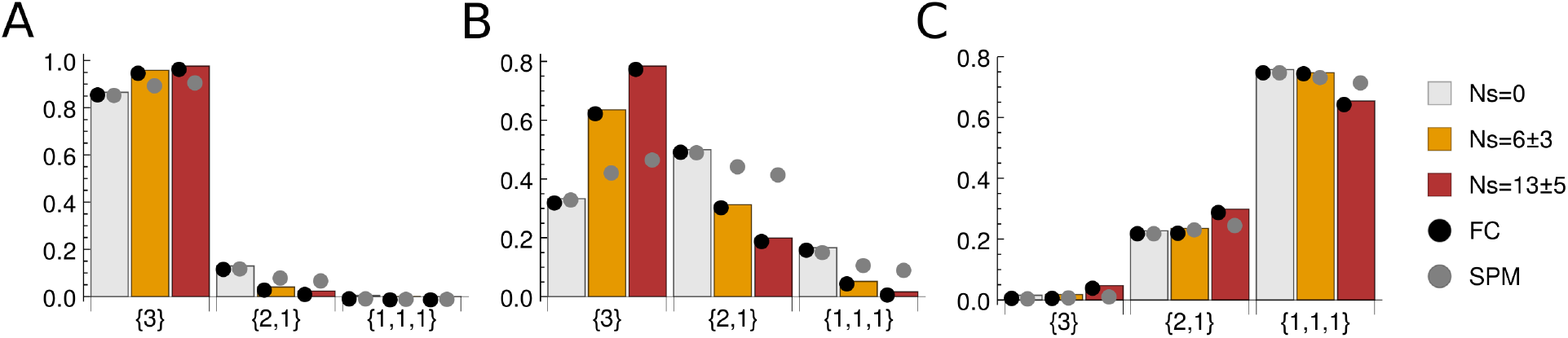
Partition probabilities for the three-plane fitness landscape. All parameters and symbols are as in Fig. 4, unless indicated otherwise; KL divergences are listed in Table 1.

### Network size effects

Although our approach is valid for an arbitrary number of alleles *K*, statistics of allele diversity in a population under selection become substantially easier to deal with in the infinite-allele limit. As discussed in the Introduction, this limit is justified since our focus here is on evolution of protein, RNA and DNA sequences, where the number of alleles grows exponentially with sequence length. Nonetheless, we have systematically investigated the extent of deviations between our infinite-allele theory results and simulations as the number of alleles *K* decreases and becomes comparable to the population size *N*. Fig. 6 shows the KL divergence between partition probabilities derived theoretically for the two-plane landscape in the infinite-allele limit (Eq. 25) and obtained numerically on finite-size FC networks. We consider three regimes: monomorphic (*Nμ* = 0.1), weakly polymorphic (*Nμ* = 1.0), and strongly polymorphic (*Nμ* = 10.0). In the latter two cases, noticeable deviations between theory and simulations begin to appear below the *K* ~ *N* regime; the agreement improves as the population becomes more monomorphic. We conclude that our theory is applicable over a wide range of mutation rates, as long as the network size is comparable to, or greater than, the population size.

**Figure 6:**
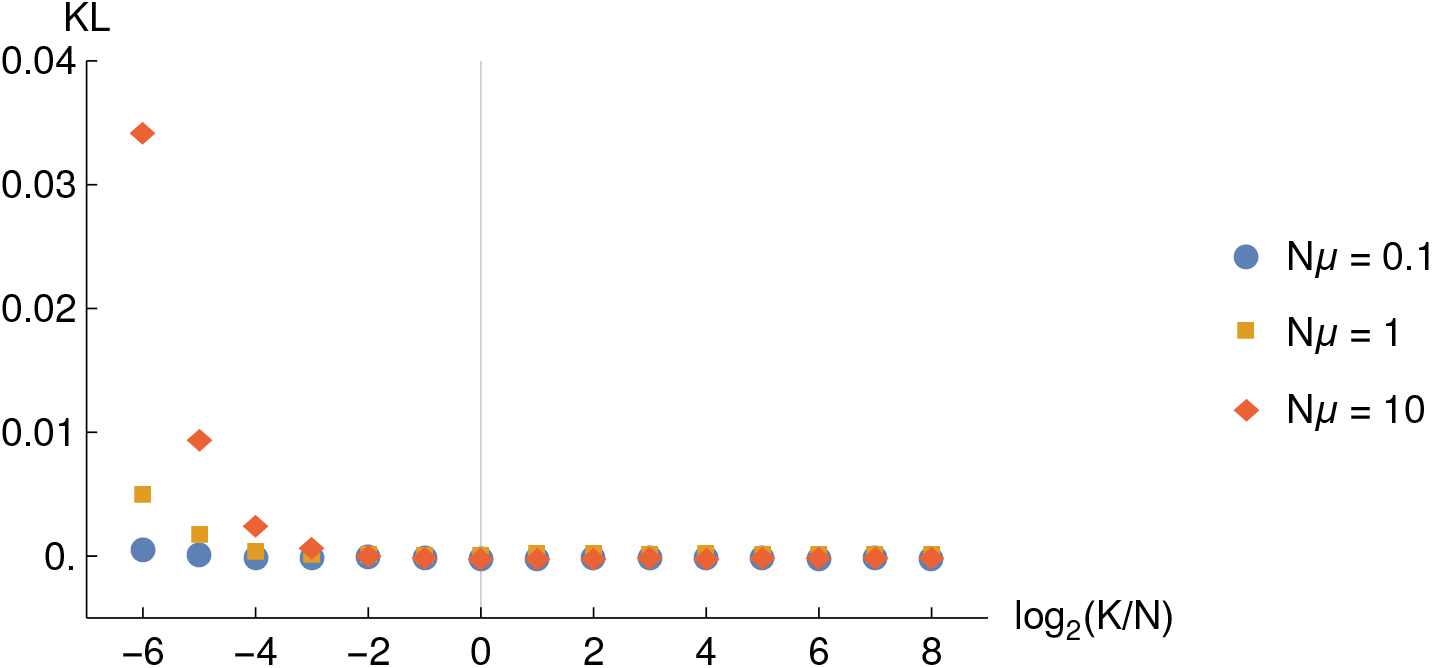
KL divergence between computational and theoretical partition probabilities on the FC two-plane fitness landscape 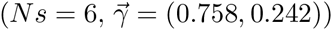, as a function of the log ratio between the total number of alleles *K* and the population size *N*. The sample size is *n* = 3; partition probabilities were estimated from 10^6^ samples as described in the main text. Population size is *N* = 10^3^, and the total number of alleles is *K* = 10^3^ × 2^*i*^, *i* ∈ {−6…8}. For smaller networks, the number of the nodes in the upper and lower planes had to be rounded to the nearest integer. Diamonds: polymorphic population (*Nμ* = 10.0), squares: weakly polymorphic population (*Nμ* = 1.0), circles: monomorphic population (*Nμ* = 0.1). The solid vertical line indicates the case of the network size equal to the population size (*K* = *N*).

**Table 1:** KL divergences between theoretical predictions and numerical simulations for the two-plane fitness landscape (Fig. 4) and the three-plane fitness landscape (Fig. 5)).

|  |  | Two-plane landscape |  |  | Three-plane landscape |  |  |
| --- | --- | --- | --- | --- | --- | --- | --- |
| | | $Ns = 0$ | $Ns = 6$ | $Ns = 13$ | $Ns = 0$ | $Ns = 6 \pm 3$ | $Ns = 13 \pm 5$ |
| $N\mu = 0.1$ | **FC** | $7 \times 10^{-7}$ | $3 \times 10^{-5}$ | $8 \times 10^{-5}$ | $1 \times 10^{-5}$ | $2 \times 10^{-6}$ | $7 \times 10^{-8}$ |
| | **SPM** | $4 \times 10^{-5}$ | $8 \times 10^{-3}$ | $2 \times 10^{-2}$ | $2 \times 10^{-6}$ | $2 \times 10^{-2}$ | $3 \times 10^{-2}$ |
| $N\mu = 1$ | **FC** | $5 \times 10^{-5}$ | $2 \times 10^{-5}$ | $1 \times 10^{-4}$ | $3 \times 10^{-5}$ | $4 \times 10^{-5}$ | $5 \times 10^{-6}$ |
| | **SPM** | $4 \times 10^{-5}$ | $2 \times 10^{-2}$ | $8 \times 10^{-2}$ | $2 \times 10^{-4}$ | $9 \times 10^{-2}$ | $2 \times 10^{-1}$ |
| $N\mu = 10$ | **FC** | $9 \times 10^{-6}$ | $8 \times 10^{-6}$ | $2 \times 10^{-5}$ | $7 \times 10^{-6}$ | $1 \times 10^{-4}$ | $2 \times 10^{-5}$ |
| | **SPM** | $4 \times 10^{-5}$ | $2 \times 10^{-4}$ | $3 \times 10^{-3}$ | $6 \times 10^{-6}$ | $1 \times 10^{-4}$ | $2 \times 10^{-2}$ |

## Discussion

One of the most challenging problems in evolutionary biology is to understand evolutionary dynamics of molecular loci, such as protein or RNA-coding sequences, or gene regulatory regions. The number of nucleotides at these loci, *L*, is large enough so that the total number of possible sequences, *K* = *A^L^*, is astronomical, far exceeding the population size *N*. Under these conditions the evolution of a molecular locus, assumed to be decoupled by recombination from the rest of the genome, reaches a “de-labelled” steady state characterized by mutation-selection-drift balance. The allelic diversity in the population is determined by the balance of forces of selection and drift on one hand, and mutation on the other. The former act to reduce allelic diversity, while the latter acts to increase it. As a result, population statistics such as the mean number of distinct alleles, or the probability of seeing a certain allelic configuration in a sample, do not change with time, even though new genotypes continue to be explored on the effectively infinite allelic network.

The neutral allelic diversity in such a system (that is, when all alleles have the same fitness) was explored by Ewens.^11,12^ The main result of that study, the Ewens sampling formula, is widely used in population genetics. The neutral landscape is a single plane, with each allele connected to the other *K* − 1 alleles. However, recent high-throughput studies connecting protein sequences with phenotypes reveal a more complex picture: generally, a functional protein such as an enzyme can be disrupted by a subset of mutations at each of its sites (e.g., through substitution of a hydrophobic residue for a hydrophilic one in the protein core). Other mutations do not significantly change protein stability, binding affinity or specificity, and are therefore effectively neutral. Occasionally, a mutation is found which increases the protein’s fitness, but these mutations are generally infrequent. Overall, recent experimental studies point to fitness landscapes comprised of multiple interconnected planes. The simplest landscape of this kind has just two distinct fitness states, with functional sequences on the upper plane and non-functional sequences on the lower plane.^1^ Multiple-plane fitness landscapes are characterized by extensive epistasis, which is likely to be pervasive in molecular evolution.^2,3,4,5^

Since molecular evolution may be described by steady-state dynamics on multiple-plane fitness landscapes, it is of great interest to generalize the Ewens sampling formula to arbitrary fitness landscapes, and to the multiple-plane class of landscapes in particular. Tractable expressions for sampling probabilities would enable inference of selection coefficients, relative plane sizes, and mutation rates, using DNA, RNA or protein sequences sampled from the population as input. Here we report an extension of the Ewens sampling formula to arbitrary fitness landscapes, focusing especially on the multiple-plane case which yields substantial simplifications in the infinite allele limit. Unlike current state-of-the-art techniques based on the Poisson random field framework,^27^ such as the sampling probability formulas developed by Desai et al.,^26^ our approach is capable of treating epistasis. However, the essential drawback of the Ewens sampling formula and its generalizations is the “fully-connected” assumption (i.e., that each allele can mutate into every other allele). Moreover, the sampling formula becomes intractable for large sample sizes, due to a large number of terms to sum over.

Therefore, in order to study the limits of applicability of our theory, we have carried out extensive comparisons with numerical simulations on multiple-plane fitness landscapes. First, we checked the full-connectivity assumption inherent in the Ewens approach by comparing the sampling probabilities of our theory with those obtained by simulation of steady-state populations evolving on single-point-mutant networks. We find that the agreement, although dependent on the details of the fitness landscape model, the values of selection coefficients, and mutation rates (and least reliable in the weakly polymorphic regime), remains strong enough overall to encourage application of our theoretical results to sequence data. Note that our model of the fitness landscape was constructed specifically to create a non-trivial distribution of neutral, deleterious and beneficial single-point mutations for the alleles, making it in some sense as distant from the fully connected network as possible. Thus we expect the deviations to be smaller (or at least not much worse) in natural systems. Second, we have checked the infinite-allele assumption by systematically reducing the number of alleles until it became lower than the population size. We find that, over a wide range of mutation rates, deviations between theory and simulations become significant only when the number of alleles approaches the population size from above. Thus our assumption of the infinite network size is justified for all loci that are long enough, such as those encoding transcribed or regulatory regions.

Robust inference of selection coefficients and mutation rates on the basis of a sample of population allelic states requires statistics of allelic diversity to deviate substantially from both the neutral expectation and the effective population size (EPS) approximation. Clearly, no inference of selection signatures is possible on the basis of limited sample data if population dynamics is close to neutral. On the other hand, in the EPS limit only the relative size of the highest-fitness plane can be inferred. By scanning over a wide range of selection coefficients and mutation rates on a two-plane fitness landscape, we have found that, although regions of neutral and EPS dynamics are roughly complementary, there are areas of parameter space characterized by deviations from both. Thus the use of our generalized Ewens sampling formula, which is valid throughout the entire parameter space, is necessary for inferring selection signatures from data. Moreover, allelic diversity generated by steady-state evolutionary dynamics on a three-plane fitness landscape is sufficiently distinct from its two-plane counterpart in the strong-selection, weakly polymorphic regime, opening up a possibility of inferring multiple selection coefficients from the data. Another hallmark of non-neutral population dynamics is de-localization of the population to multiple fitness planes. With a two-plane landscape, we expect the fraction of the population on the lower plane to increase with the mutation rate and decrease with the distance between the two planes. Our investigation of the mutation load confirms these predictions.

In summary, we have generalized the Ewens sampling formula to evolutionary dynamics under selection. Although in principle our results are valid for arbitrary fitness landscapes, focusing on the infinite allele limit and landscapes with just two or three distinct fitness states yields substantial simplifications, making our approach computationally tractable and thus applicable to inferring selection signatures from high-throughput sequence data. Such multiple-plane fitness landscapes are consistent with recent large-scale studies of molecular phenotypes.^1,3,6,7^ Unlike previous approaches, we do not assume the absence of epistasis, which is likely to be prevalent in molecular evolution.^2,3,4,5^ However, we do make the infinite allele assumption, and, as in the Ewens original formula,^12^ assume that each allele can mutate into any other allele. We check our theory against numerical simulations in model systems where these assumptions are relaxed, and find that our predictions remain accurate enough to enable inference of evolutionary parameters from sequencing data.

## Acknowledgements

PK acknowledges financial support from a research fellowship awarded by the Department of Physics and Astronomy, Rutgers University. AVM was supported in part through a collaboration with Los Alamos National Lab (LANL-DOE 20150236ER).

## Data Availability

Software and models used in this study are freely available upon request.

## Appendix

## A Simplification of the sampling formula in the two-plane system

In the two-plane system, the fitness vector has the following structure:

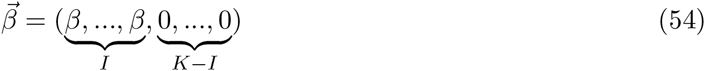

with *I* nonzero entries followed by *K* − *I* zeros. In this case, Eq. 6 involves summation over only *I* indices:

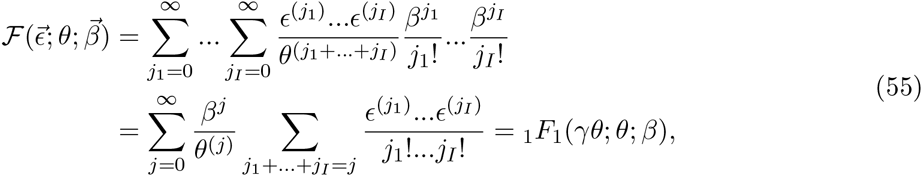

where *γ* = *I*/*K* is the fraction of nodes on the first (lower) plane, and in the last equality we used

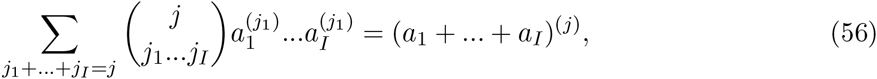

where the sum runs over all non-negative *j_i_* that sum up to *j*.

Now, consider a situation in which the first *i* out of *k* counts happen to come from the first plane. This means that they are among the first *I* elements of 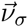. Since assigning these *i* counts to different locations within the first *I* slots in the 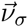 vector (while keeping their original order) does not change the result, we can for convenience assign them to be the first *i* elements of 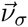, followed by *I* − *i* zeros. Then

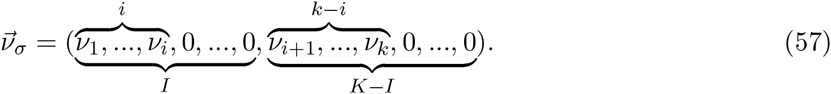

The corresponding generalized confluent hypergeometric function is again given by the sum over *I* indices:

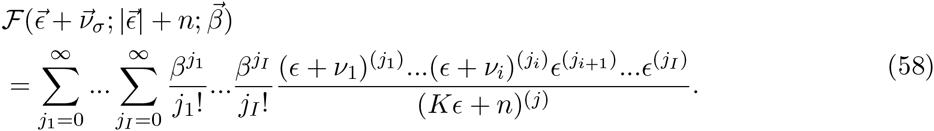

We can rewrite it as

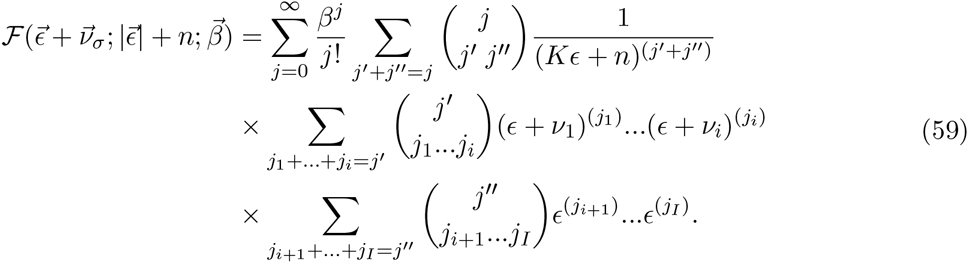

Using Eq. 56 immediately leads to

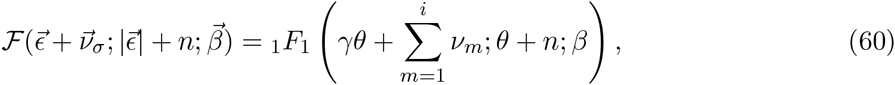

so that the generalized confluent hypergeometric function is once again reduced to the ordinary confluent hypergeometric function.

Lastly, we need to take into account the fact that we can put counts into different positions of the 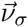 vector. This introduces an additional binomial pre-factor 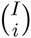. Similarly, placing the rest of the counts into the last *K* − *I* entries of the 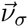 vector introduces another binomial pre-factor 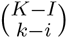. Using these pre-factors together with Eqs. 55 and 60 in Eq. 20 yields Eq. 24.

## B Frequency spectrum for the arbitrary landscape

We can expand the exponents in the allele frequency distribution (Eq. 4) into a series:

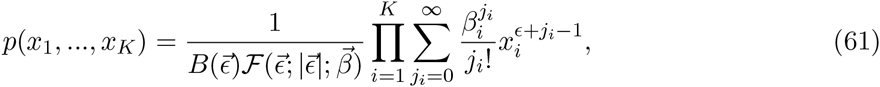

and apply

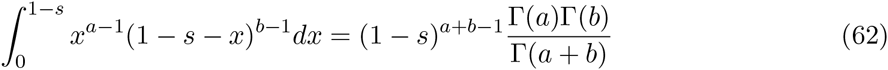

in order to get

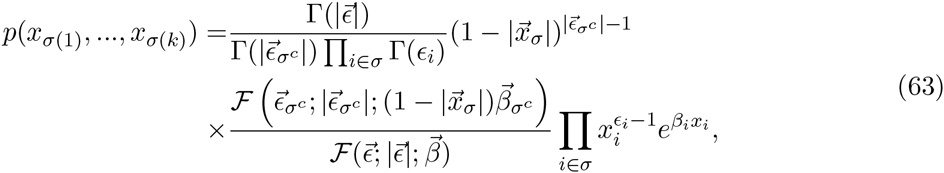

where *σ^c^* is a list of the *K* − *k* alleles not contained in *σ*, and therefore 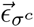 and 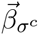 are (*K* − *k*)-dimensional vectors obtained from 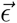 and 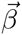 by eliminating elements at the positions specified by *σ*, while 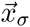 is a *k*-dimensional vector obtained from the *K*-dimensional vector 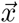 by keeping the elements at the positions specified by *σ*, and eliminating the rest.

With equal mutation rates, we have

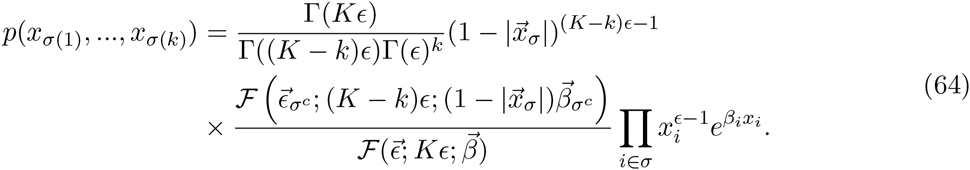

Taking the *k* = 1 case and summing over allelic types, we obtain Eq. 34.

## Footnotes

a All the formulas in this section can be generalized to the case of final-state-dependent mutation rates, i.e. *μ_ij_* = *μ_j_*, ∀*i* in *A_i_* → *A_j_*

b We shall use the notation {*a, b*, …, *z*} for unordered lists and (*a, b*, …, *z*) for ordered ones. For ordered lists (*a, b*, …, *z*) ≠ (*b, a*, …, *z*), whereas for unordered lists {*a, b*, …, *z*} = {*b, a*, …, *z*}.

c We shall use the notation (*a*_1_, *a*_2_, …, *a_n_*) for allelic histograms.

## References

[1] Podgornaia, A. and Laub, M. (2015) Science 347, 673–677.

[2] Lunzer, M., Miller, S. P., Felsheim, R., and Dean, A. M. (2005) Science 310, 499–501.

[3] Romero, P. A. and Arnold, F. H. (2009) Nat Rev Mol Cell Biol 10, 866–876.

[4] Lunzer, M., Golding, G. B., and Dean, A. M. (2010) PLoS Genet 6, e1001162.

[5] Breen, M., Kemena, C., Vlasov, P., Notredame, C., and Kondrashov, F. (2012) Nature 490, 535–538.

[6] Lind, P. A., Berg, O. G., and Andersson, D. I. (2010) Science 330, 825–827.

[7] Hietpas, R. T., Jensen, J. D., and Bolon, D. N. A. (2011) Proc Nat Acad Sci USA 108, 7896–7901.

[8] Sanjuan, R., Moya, A., and Elena, S. F. (2004) Proc Nat Acad Sci USA 101, 8396–8401.

[9] Eyre-Walker, A. and Keightley, P. D. (2007) Nat Rev Genet 8, 610–618.

[10] Wagner, A. (2008) Nat Rev Genet 9, 965–974.

[11] Ewens, W. (2004) Mathematical Population Genetics: I. Theoretical Introduction, Springer, 2nd edition.

[12] Ewens, W. (1972) Theor Pop Biol 3, 87–112.

[13] Slatkin, M. (1994) Genet Res Cambr 64, 71–74.

[14] Li, W.-H. (1978) Genetics 90, 349–382.

[15] Li, W.-H. (1977) Proc Nat Acad Sci USA 74, 2509–2513.

[16] Li, W.-H. (1979) Genetics 92, 647–667.

[17] Ewens, W. and Li, W.-H. (1980) J Math Biol 10, 155–166.

[18] Griffiths, R. (1983) Journal of Mathematical Biology 17, 1–10.

[19] Ethier, S. and Kurtz, T. (1987) Stochastic Models in Biology, Lecture Notes in Biomathematics 70, 72–86.

[20] Joyce, P. and Tavare, S. (1995) J Math Biol 33, 602–618.

[21] Joyce, P. (1995) J Appl Prob 32(3), 609–622.

[22] Grote, M. and Speed, T. (2002) Ann Appl Prob 12, 637–663.

[23] Joyce, P., Genz, A., and Buzbas, E. (2012) J Comp Biol 16(6), 650–661.

[24] Charlesworth, B., Morgan, M., and Charlesworth, D. (1993) Genetics 134, 1289–1303.

[25] Hudson, R. and Kaplan, N. (1994) Gene trees with background selection In B. Golding, (ed.), Non-Neutral Evolution: Theories and Molecular Data, pp. 140–153 Chapman and Hall New York, NY.

[26] Desai, M., Nicolaisen, L., Walczak, A., and Plotkin, J. (2012) Theor Pop Biol 81, 144–157.

[27] Sawyer, S. and Hartl, D. (1992) Genetics 132, 1161–1176.

[28] Moran, P. A. P. (1958) Math Proc Cambr Philos Soc 54, 60–71.

[29] Gillespie, J. (2004) Population Genetics: A Concise Guide, The Johns Hopkins University Press, Baltimore.

[30] Wright, S. (1931) Genetics 16, 97–159.

[31] Mustonen, V. and Lassig, M. (2010) Proc Nat Acad Sci USA 107(9), 4248–4243.

[32] Watterson, G. (1977) Genetics 85, 789–814.

[33] Kimura, M. (1962) Genetics 47, 713–719.

[34] Kimura, M. and Ohta, T. (1969) Genetics 61, 763–771.

[35] Crow, J. and Kimura, M. (1970) An Introduction to Population Genetics Theory, The Blackburn Press, Caldwell, NJ.

[36] Sella, G. and Hirsh, A. (2005) Proc Nat Acad Sci USA 102, 9541–9546.

[37] Sella, G. (2009) Theor Pop Biol 75, 30–34.

[38] Rouzine, I. M., Rodrigo, A., and Coffin, J. M. (2001) Microbiol Mol Biol Rev 65, 151–185.

[39] Kingman, J. F. C. (1975) Journal of the Royal Statistical Society, B 37(1), 1–22.

[40] Kingman, J. F. C. (1977) Theor Pop Biol 11(2), 274–283.

[41] Etheridge, A. (2011) Some Mathematical Models from Population Genetics, Springer-Verlag, Berlin 1st edition.

[42] Nielsen, R. (2005) Annu Rev Genet 39, 197–218.

[43] Kullback, S. and Leibler, R. (1951) Ann Math Stat 22, 79–86.

